# A putative enoyl-CoA hydratase contributes to biofilm formation and the antibiotic tolerance of *Achromobacter xylosoxidans*

**DOI:** 10.1101/620559

**Authors:** Lydia C. Cameron, Benjamin Bonis, Chi Q. Phan, Leslie A. Kent, Alysha K. Lee, Ryan C. Hunter

## Abstract

*Achromobacter xylosoxidans* has attracted increasing attention as an emerging pathogen in patients with cystic fibrosis. Intrinsic resistance to several classes of antimicrobials and the ability to form robust biofilms *in vivo* contribute to the clinical manifestations of persistent *A. xylosoxidans* infection. Still, much of *A. xylosoxidans* biofilm formation remains uncharacterized due to the scarcity of existing genetic tools. Here we demonstrate a promising genetic system for use in *A. xylosoxidans*; generating a transposon mutant library which was then used to identify genes involved in biofilm development *in vitro*. We further described the effects of one of the genes found in the mutagenesis screen, encoding a putative enoyl-CoA hydratase, on biofilm structure and tolerance to antimicrobials. Through additional analysis, we find that a cis-2 fatty acid signaling compound is essential to *A. xylosoxidans* biofilm ultrastructure and maintenance. This work describes methods for the genetic manipulation of *A. xylosoxidans* and demonstrated their use to improve our understanding of *A. xylosoxidans* pathophysiology.

## INTRODUCTION

Cystic fibrosis (CF) is caused by mutations in the gene encoding the cystic fibrosis transmembrane conductance regulator (CFTR) protein that resides at the apical surface of many epithelial cell types. CFTR defects result in abnormal chloride and bicarbonate transport, increasing mucus viscosity in the pancreas, paranasal sinuses, digestive tract, and most notably, the lower airways. Viscous mucus, due to its impaired clearance and nutrient bioavailability, in turn becomes chronically colonized by pathogenic bacteria that are the leading cause of mortality among the CF population (1). For decades, *Pseudomonas aeruginosa* and *Staphylococcus aureus* have been extensively characterized and recognized as the primary airway pathogens. More recently, however, many other multidrug-resistant opportunistic bacterial species have received attention for their association with CF disease progression (2).

Among them, *Achromobacter xylosoxidans* is notable given its association with poor pulmonary function scores and apparent patient-to-patient transmissibility (3-5). This aerobic, Gram-negative opportunistic pathogen has been found to colonize anywhere from 2 to 20 percent of CF subjects, though its prevalence has risen in recent years (4, 6-11), sparking a renewed interest in its pathophysiology. Of particular concern is the intrinsic and acquired resistance of *A. xlyosoxidans* to multiple classes of antimicrobial agents, including aminoglycosides, beta-lactams, carbapenems, chloramphenicol and fluoroquinolones, which presents a significant burden for infection control (7, 8, 12-15).

Also complicating treatment of *A. xylosoxidans* is its ability to form robust biofilms (16-18). Thought to be the predominant *in vivo* lifestyle among CF pathogens (19), biofilms confer a highly protective growth environment that shields pathogens against environmental stress, antimicrobials and the host immune response (20). In addition, nutrient and oxygen gradients throughout microcolonies (or aggregates)(16) result in slowed metabolism among biofilm cells which can also potentiate drug tolerance. The recalcitrance of biofilm cells to antibiotic exposure likely enhances the persistence of *A. xylosoxidans* during chronic infection of the CF lung.

Relative to canonical CF pathogens, comparatively little is known about the molecular mechanisms of biofilm formation and maintenance in *A. xylosoxidans*. This is due, in part, to the paucity of genetic tools available for manipulation of its genome. Therefore, the objectives of this study were two-fold: First, we sought to develop a tractable system to genetically manipulate *A. xylosoxidans*. Using this system, our second objective was to take a transposon mutagenesis approach to study the molecular basis of *A. xylosoxidans* biofilm formation. In doing so, we identified several gene products essential for biofilm development. We chose to further characterize a gene identified most frequently in our screen (*echA)* encoding a putative enoyl-CoA hydratase that, when disrupted, leads to a decrease in biofilm accumulation and increased susceptibility to multiple classes of antibiotics.

## RESULTS

### Identification of biofilm-defective mutants via transposon mutagenesis

To identify genetic determinants of biofilm formation in *A. xylosoxidans*, we generated a mutant library of strain MN001 using random transposon mutagenesis employing the mini-mariner transposable element (21). From this library, 15,000 mutants were screened for altered biofilm formation using an established crystal violet (CV) / microtiter dish assay (22). After an initial round of screening, 134 putative biofilm-defective mutants were identified, and putative hits were then re-tested (n=4) to verify mutant phenotypes. After secondary screening and eliminating mutants with general growth defects, 31 transposon mutants were confirmed to be defective in biofilm accumulation (Fig. 1, Table 1).

**Table 1.** List of independent biofilm-defective transposon mutants identified in this study.

| Mutant | Axylo Locus Tag | Annotation | Insertion Coordinate |
| --- | --- | --- | --- |
| 18-C10 | 2959 | Extracellular serine protease precursor | 3300783 |
| 21-A5 | 0405 | Enoyl-CoA hydratase | 458026 |
| 21-B10 | 2974 | DoxX family protein | 3318225 |
| 21-F3 | 0405 | Enoyl-CoA hydratase | 458968 |
| 37-D9 |  | No sequence |  |
| 29-D11 | 1017 | Branched chain amino acid transport protein | 1140377 |
| 29-F4 | 3436 | Flagellar basal body rod protein FlgB | 3820439 |
| 34-H5 | 1568 | Carboxymethylenebutenolidase | 1767696 |
| 44-D4 |  | No sequence |  |
| 45-B11 | 1417 | Outer membrane protein assembly factor BamD | 1595377 |
| 52-D3 | 2959 | Extracellular serine protease precursor | 3296785 |
| 56-D9 |  | No sequence |  |
| 61-A2 | 4754 | Hypothetical protein | 5265710 |
| 61-G3 | 0328 | Hypothetical protein | 374132 |
| 68-D1 | 2959 | Extracellular serine protease precursor | 3303694 |
| 75-D6 | 0867 | Propionate hydroxylase | 961203 |
| 77-F11 | 2959 | Extracellular serine protease precursor | 3302134 |
| 79-B9 | 0165 | Glycosyltransferase CsbB | 200048 |
| 83-E10 | 0126 | Hypothetical protein | 152964 |
| 84-D4 | 0867 | Propionate hydroxylase | 961404 |
| 86-E10 | 0405 | Enoyl-CoA hydratase | 458180 |
| 87-B2 | 3872 | Glycosyltransferase MshA | 4315184 |
| 90-D7 |  | No sequence |  |
| 95-D11 | 1003 | Hypothetical protein | 1124527 |
| 103-G8 | 5298 | L-asparaginase | 5841569 |
| 114-B2 | 0867 | Propionate hydroxylase | 961364 |
| 114-E11 |  | No sequence |  |
| 115-C6 | 0951 | LysR family transcriptional regulator DmlR | 1065410 |

**Figure 1.**
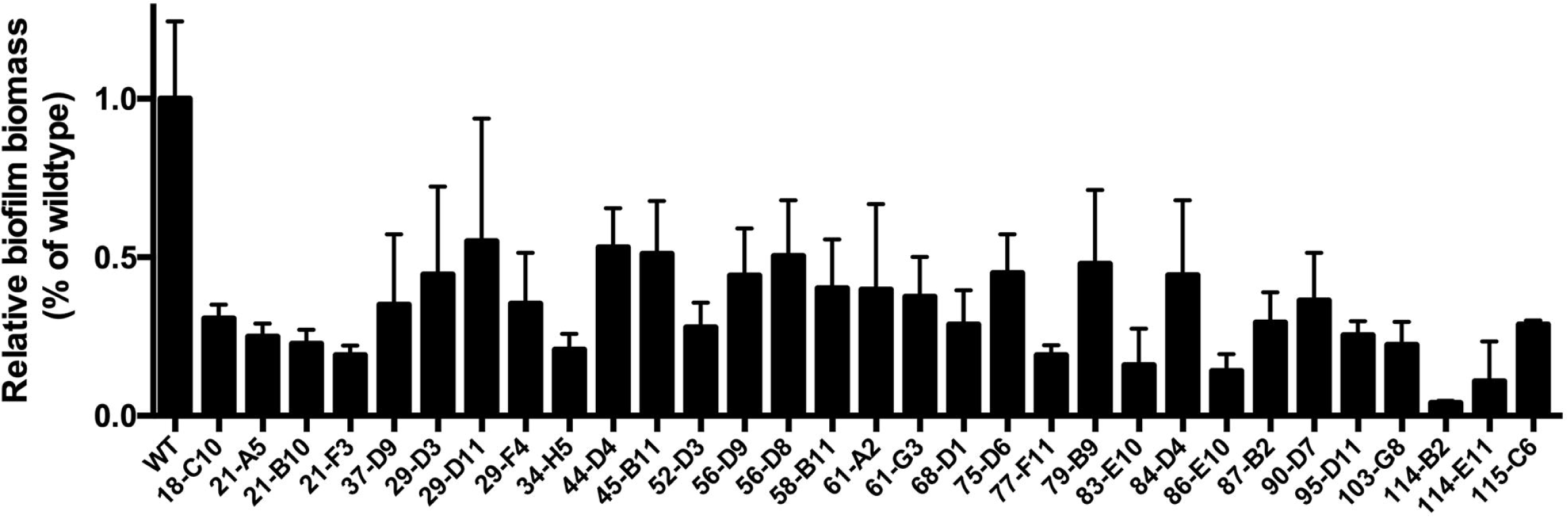
Biofilm formation by transposon mutants of *A. xylosoxidans* MN001. Mean crystal violate absorbance for each mutant is expressed relative to the parental wildtype strain. Error bars represent standard deviation of the mean (n=4).

To identify DNA sequences (∼200-400 bp) flanking transposon insertion junctions, mutants were screened by arbitrarily-primed PCR. Insertions were mapped to loci predicted to encode various classes of gene products, including membrane proteins, glycosyltransferases, flagella, transcriptional regulators, in addition to several proteins with no assigned function (Table 1). Arbitrary PCR of three transposon insertion loci generated sequences with no homology to published gene sequences, and we were unable to obtain transposon-flanking sequences for five additional mutants. Of note, a putative enoyl-CoA hydratase, an exonuclease, and a propionate hydroxylase were each hit multiple times. We took interest in the gene encoding a putative enoyl-CoA hydratase (Axylo_0405) given the importance of homologous proteins in fatty acid signal biosynthesis and biofilm development among diverse bacterial species (23-29). We named this gene *echA* (for <u>e</u>noyl-<u>C</u>oA <u>h</u>ydratase <u>A</u>) and sought to characterize its effects on *A. xylosoxidans* biofilm physiology in greater detail.

### Generation and complementation of an *echA* mutant

Genetic tractability has been challenging for *A. xylosoxidans* due its inherent resistance to multiple classes of antibiotics and inducible efflux system as a second line of defense (12,13,15). We therefore sought to develop a robust genetic system for the generation and complementation of marker-free deletion mutants that would facilitate further study of the genetic basis of biofilm formation. Given that tetracycline resistance was an effective marker to generate the transposon mutant library, the tetracycline resistance cassette from pEX18tc (30) was PCR amplified and ligated into the mobilizable suicide vector pSMV8 (31) generating pBMB1 (empty vector) and pBMB2 (*echA* deletion construct; Figure S1). Attempts to construct a complementation vector using broad host-range cloning plasmids were unsuccessful. We therefore introduced *echA* into pBMB3 (pBBR1MCS-5::*alcA*) forming pBMB4 (Figure S1), placing *echA* expression under the control of an alcohol-inducible promoter (32). Using these vectors, we generated a markerless deletion mutant and complement of *A. xylosoxidans* MN001 lacking *echA*.

### *A. xylosoxidans* biofilm ultrastructure is mediated by *echA*

To confirm that the biofilm-deficient phenotypes of transposon mutants 21-A5, 21-F3, and 86-E10 (Table 1) were a result of mutations in *echA*, we re-assessed biofilm formation by the markerless Δ*echA* mutant using the CV assay described above. As before, Δ*echA* exhibited a significant decrease in biofilm biomass accumulation relative to the wildtype (p=0.002) after 72h (Fig. 2). Complementation with pBMB4 restored the wildtype phenotype (p=0.023), confirming a role of *echA* and its gene product in *A. xylosoxidans* biofilm development.

**Figure 2.**
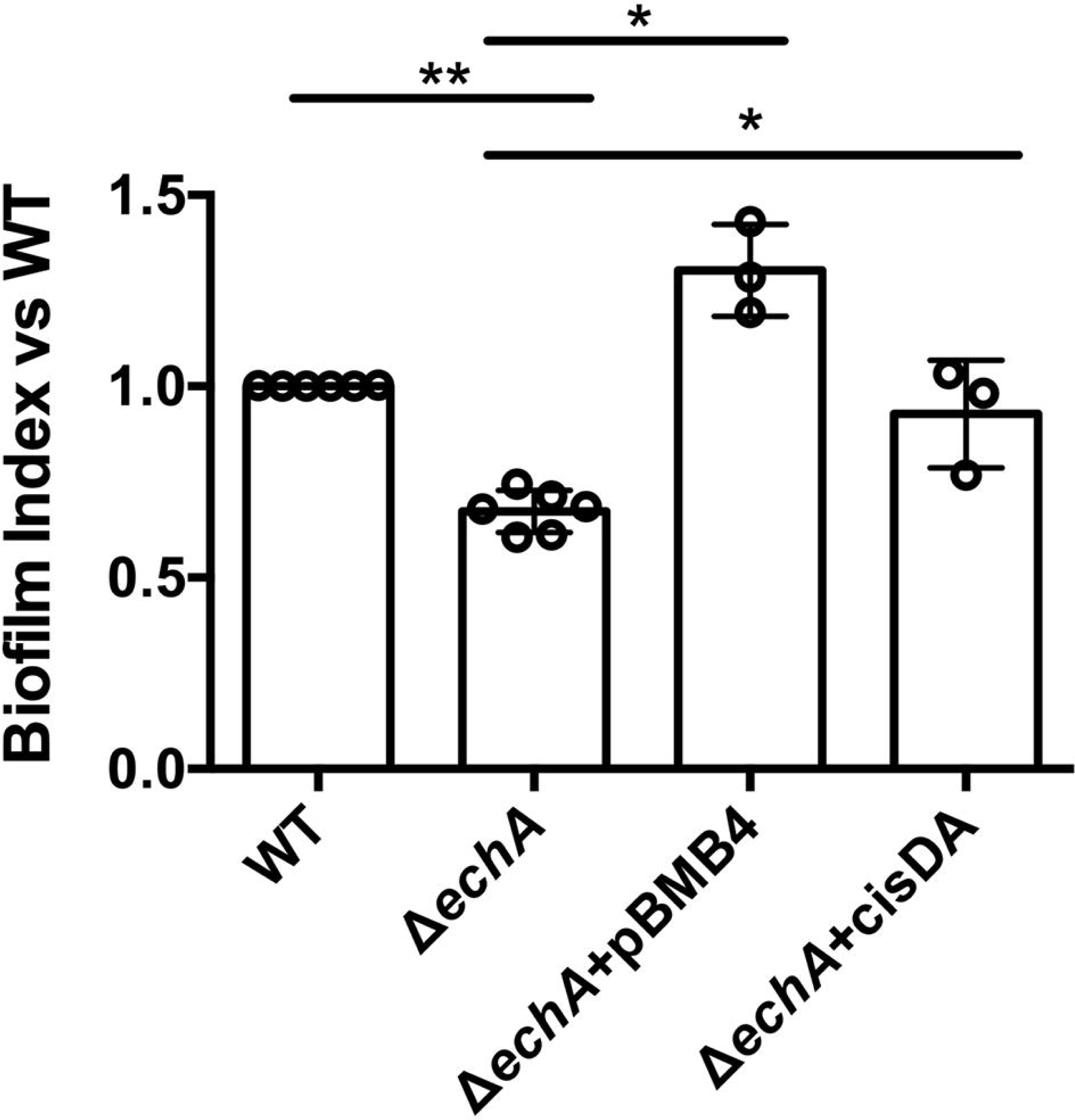
A putative enoyl-CoA hydratase contributes to *A. xylosoxidans* biofilm formation via biosynthesis of a fatty acid signaling metabolite. *A. xylosoxidans* biofilms were grown for 72 h, quantified using crystal violet and normalized to cell density (biofilm index). Deletion of *echA* resulted in a significant decrease in biofilm biomass that was restored via complementation and addition of cis-DA. Error bars represent standard error of the mean (n=6 for WT and Δ*echA*; n=3 for Δ*echA*+pBMB4 and Δ*echA*+cisDA).

Enoyl-CoA hydratases are central to the biosynthesis of a class of fatty acid signaling molecules, or diffusible signal factors (DSFs), that have been described in diverse bacterial species for their role in mediating virulence, motility and biofilm development (24-29, 33, 34). In *P. aeruginosa,* for example, the DSF *cis*-2-decenoic acid (cis-DA) is known to play a critical role in biofilm dispersion (24,35). Therefore, to test whether the observed biofilm impairment phenotype in *A. xylosoxidans* was also mediated by a DSF-like signaling mechanism, we exogenously added synthetic cis-DA (310nM)(35) to the Δ*echA* mutant during biofilm development. Despite having no effect on the WT strain (not shown), addition of cis-DA to Δ*echA* resulted in a significant restoration of biofilm biomass relative to the untreated control (p=0.024)(Fig. 2). These data suggest that biofilm formation in MN001 is directly mediated by a DSF-like metabolite generated by an enoyl-CoA hydratase.

Since the CV staining approach used in the initial mutant screen relies on a dye that stains not only cells, but all biomass adhering to the microtiter plate, we elected to use additional biofilm assays to further characterize the biofilm phenotype of Δ*echA,* and whether disruption of the putative enoyl-CoA hydratase negatively impacts a specific stage of biofilm development (*e.g.* attachment, matrix production, maturation). We first used a modified adhesion assay (36) to test for defects in attachment to the surface of a chamber slide. Under static conditions, Δ*echA* exhibited no difference in surface attachment relative to the wildtype, demonstrating that initial stages of biofilm formation are not affected (Fig. 3A). We then used a common colony morphology assay that relies on Congo red staining as a macroscopic means of identifying the capacity for extracellular matrix production (37,38). Over the course of 6 days of growth, biofilm colonies showed no apparent differences in morphology or color between strains, suggesting that the *echA* gene product has negligible effect on matrix production in *A. xylosoxidans.* Finally, we used scanning electron microscopy to visualize microcolony ultrastructure in mature biofilms grown for 72 hours under static conditions (Fig. 3C). Relative to WT, the Δ*echA* mutant exhibited notable phenotypic differences in biofilm architecture, corroborating observations made using the microtiter plate assay. First, the mutant strain demonstrated less robust biofilm growth and reduced surface coverage relative to WT. In addition, higher magnification images revealed a less-dense packing order to mutant biofilm cells, suggesting that cell-cell signaling mediated by enoyl-CoA hydratase-derived metabolite is central to biofilm ultrastructural development in *A. xylosoxidans.*

**Figure 3.**
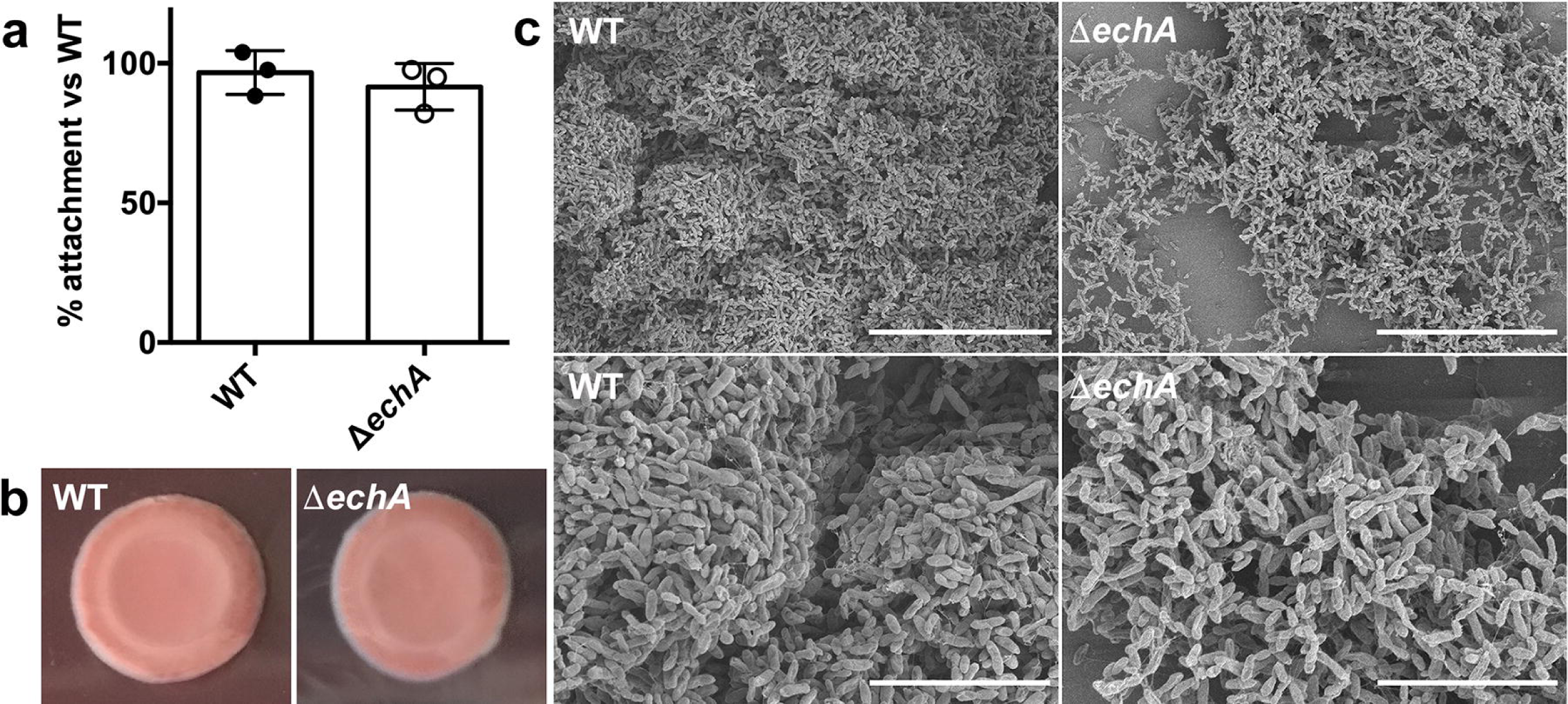
echA plays a central role in biofilm ultrastructure in *A. xylosoxidans* MN001. (a) Early attachment assay for WT and *echA* mutant. Error bars represent standard deviation of the mean for replicate experiments (n=3). (b) Colony biofilms grown on Congo red plates for 6 days revealed no differences in matrix production. (c) Mature biofilms of the WT and mutant were visualized by SEM to examine biofilm architecture (scale bars, top = 30μm, bottom = 10 μm). All images shown are representative of experiments performed in biological triplicate (n=3) with similar results.

### Δ*echA* exhibits increased susceptibility to antibiotic treatment

Biofilm-associated bacteria can be orders of magnitude more tolerant to antibacterial compounds than their planktonic counterparts (39). Given its defect in biofilm formation and more diffuse nature of mature microcolonies shown by SEM, we hypothesized that the Δ*echA* mutant would exhibit increased susceptibility to antimicrobial treatment. To test this, mature biofilms were grown statically in 8-chamber coverslip slides for 72h and treated with various concentrations of two commonly used therapeutics to which *A. xylosoxidans* is tolerant, levofloxacin and tobramycin (Fig. 4). Despite not showing any difference in minimum inhibitory concentrations for planktonic cells of the WT and mutant strains (100 μg/mL for tobramycin; 4 μg/mL for levofloxacin), Live/Dead staining of treated biofilms, visualization by microscopy (Fig. 4A), and quantification of dead biomass (Fig. 4B) revealed a statistically significant increase (p<0.001) in susceptibility of the Δ*echA* biofilm to both antibiotics relative to WT. These data demonstrate a central role of *echA* in biofilm antibiotic tolerance.

**Figure 4.**
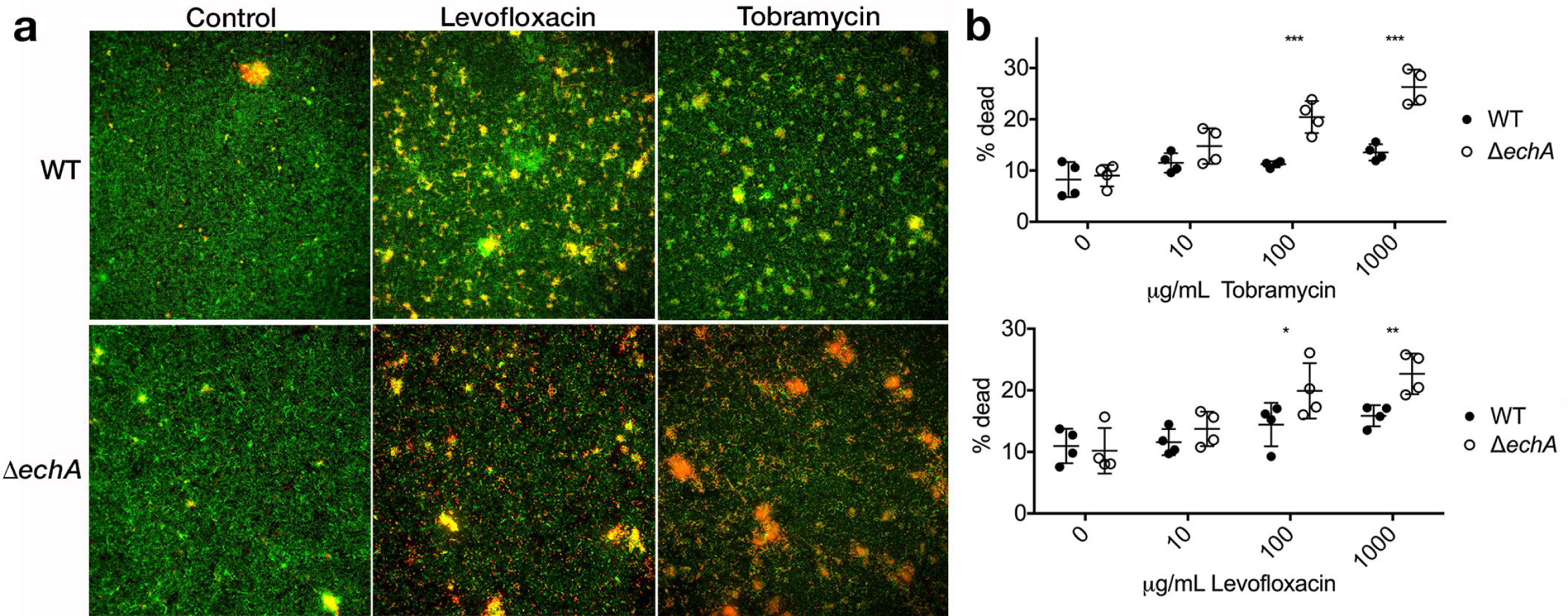
Disruption of enoyl-CoA hydratase activity leads to increased antibiotic susceptibility. (a) Mature *A. xylosoxidans* biofilms were visualized by fluorescence microscopy after 6h exposure to levofloxacin (100μg/mL) and tobramycin (1000μg/mL). Deletion of *echA* resulted in increased susceptibility to both compounds. (b) Integrative densities of green (live) and red (dead) cells were used to calculate % dead biomass for each biofilm treated with various concentrations of antibiotic (bars represent standard deviation of the mean, n=4 for each treatment).

## DISCUSSION

*A. xylosoxidans* is recognized as an emerging nosocomial pathogen and is associated with a range of infections, including bacteremia (40), endocarditis (41), meningitis (42), and pneumonia in immunocompromised individuals (43). In addition, the prevalence of *A. xylosoxidans* is now estimated to be as high as 20% among individuals with CF (6-11), which is of increasing concern given its reported correlation to lung function decline (3), patient-to-patient transmissibility (5), and multi-drug resistance phenotypes (12-15). Unfortunately, little is known about the physiology and molecular biology of this pathogen, and a greater understanding is critically needed to inform new treatment strategies. This study is an important step in that direction as it identifies several molecular determinants of biofilm growth, thought to play a critical role in the persistence and pathogenesis of *A. xylosoxidans* in the CF airways (16).

The inherent and acquired drug resistance of *Achromobacter* spp. not only limits therapeutic approaches but has also led to a paucity of tools (i.e. selection markers) available for molecular biology studies of *A. xylosoxidans* and manipulation of its genome. Here, we systematically screened numerous vectors and determined that among antibiotics tested, only tetracycline and gentamicin resistance markers were compatible with both *E. coli* and *A. xylosoxidans*. Tet^R^ and Gent^R^ cassettes were then used to generate (to our knowledge) the first transposon mutant library and a null deletion strain in *A. xylosoxidans*, facilitating a detailed study at the molecular level. These tools pave the way for addressing critical questions about the physiology of this bacterium and its role in pathogenesis.

Screening of our transposon library revealed several genetic determinants of biofilm formation. Statistically, genome coverage in our library was unsaturated (∼92%), though 31 hits in genes in diverse functional categories suggests that the regulation of biofilm development in *A.xylosoxidans* integrates a complex network of cellular processes and environmental stimuli. Several homologs of genes identified in this screen have been linked to biofilm formation in other bacteria. For example, *flgB,* encoding a flagellar basal body rod protein, has been implicated in biofilm development in both *Bordetella bronchiseptica* (44) and *P. aeruginosa* (45). It is noteworthy that other flagellar-associated genes are downregulated in *A. xylosoxidans* biofilms relative to planktonically grown cells (46). In addition, LysR family regulators, such as Achromo_0951 that was identified in our screen, have been implicated in biofilm morphology and mucosal colonization in other respiratory pathogens such as *P. aeruginosa, Burkholderia cenocepacia, and Klebsiella pneumoniae* (47-49). Further characterization of these and other genes identified in our screen (see Table 1) will undoubtedly generate new insights into the many aspects of *Achromobacter* biofilm physiology that remain uncharacterized.

We took most interest in Axylo_0405 (*echA*), encoding a putative enoyl-CoA hydratase (one of eight in strain MN001) with homology to DspI in *P. aeruginosa* (24). DspI and other bacterial homologs (e.g. RpfF in *X. campestris*)(27) are key enzymes used in the synthesis of diffusible signaling factors (DSFs) that are monounsaturated fatty acids of medium chain length containing a cis-2 double bond thought to be central in their mechanism of action. These metabolites broadly regulate bacterial behaviors such as motility, iron uptake and virulence (50-52), and data presented here adds to the growing appreciation of their widespread role in biofilm development across diverse Gram-negative bacteria. While synthetic cis-DA was able to partially rescue the biofilm defect in Δ*echA*, the specific identity of the fatty acid signal produced by *A. xylosoxidans* remains to be determined. It is also possible that *A. xylosoxidans* may respond to multiple DSF-like signals produced endogenously or exogenously, as interspecies signaling mediated by cis-2 fatty acids has also been reported (28). Ongoing work is now aimed at characterizing the signal structure, its biosynthetic pathway, sensing mechanism(s) and downstream phenotypic effects in addition to biofilm development.

Respiratory pathogens that adopt a biofilm lifestyle *in vivo* exist in a protective environment against antimicrobials and host defense mechanisms. Furthermore, biofilm cells generally reduce their metabolic activity relative to planktonic cells, further enhancing their intrinsic resistance to therapy. *A. xylosoxidans* has been shown to form robust biofilms both *in vivo* and *in vitro* (16,18), which is corroborated by microscopy data presented here. However, our data demonstrating a heightened susceptibility of Δ*echA* to antibiotic treatment suggest that interfering with fatty acid-mediated cell-cell signaling may represent a viable approach to managing *A. xylosoxidans* biofilms in the lower airways, particularly when used in combination with conventional CF antibiotics. Here we elected to test two compounds to which *A.xylosoxidans* has high resistance, levofloxacin and tobramycin (53), though it will be interesting to test whether this effect holds true for other antibiotics with different mechanisms of action.

In summary, we developed a robust genetic system for use in *A. xylosoxidans* that we leveraged to generate a transposon mutant library. Further study of this library revealed a suite of genes essential for biofilm development *in vitro* and invoked an essential role for a putative enoyl-CoA hydratase in biofilm ultrastructure and tolerance to antimicrobials, likely mediated by a cis-2 fatty acid signaling compound. While a clear picture of the clinical impact of *A. xylosoxidans* in CF airway disease is only beginning to emerge, the continued study of the molecular basis of its biofilm formation and physiology will undoubtedly lead to improved treatment strategies for this important respiratory pathogen.

## METHODS

### Bacterial strains, plasmids, and growth conditions

Bacterial strains used are listed in Table S1. *A. xylosoxidans* MN001 (54) and *Escherichia coli* were grown in lysogeny broth (LB) at 37°C unless otherwise specified. When necessary, growth media were supplemented with gentamicin at 20 μg/ml (*E. coli*) or 100 μg/ml (*A. xlyosoxidans*), ampicillin at 100 μg/ml (*E. coli*) or 300 μg/ml (*A. xylosoxidans*), carbenicillin (300 μg/ml), tetracycline (300 μg/ml), or chloramphenicol (30 μg/ml). *E. coli* strain β2155 was supplemented with 60 mM diaminopimelic acid (DAP).

### Transposon mutagenesis

The transposon delivery vector pTnTet containing the hyperactive mariner transposon (21) was introduced by transformation into the *E. coli* donor bacterial strain β2155 (55). *A. xylosoxidans* MN001 and *E. coli* β2155 (carrying pTnTet) were grown overnight in LB and LB containing chloramphenicol (30mg/ml) and 60 mM DAP, respectively, at 37°C. Cells were combined in a donor-to-recipient ratio of 5:1, centrifuged at 4000 x *g* for 5 minutes, resuspended in 200 mL fresh LB-DAP and spot-plated (10 μL) onto LB agar. Mating proceeded for 8h at 37°C, at which point cells were harvested and resuspended in 1 mL of LB. Cell suspensions were diluted 1:5, and 100 mL aliquots were plated on LB-tetracycline agar and allowed to grow for 48h at 37°C. Colonies were randomly selected for downstream biofilm assays.

### Biofilm microtiter plate assay

To identify determinants of biofilm development, transposon mutants were screened using a modified crystal violet (CV) assay approach described previously (22). Briefly, individual mutants were transferred to single wells of a 96-well microtiter plate containing 200 μL LB per well, and incubated while shaking at 37°C. Following 24h of growth, 2 μL was transferred to 198 μL ABT medium [15mM (NH_4_)_2_SO_4_, 40mM Na_2_HPO_4_, 20mM KH_2_PO_4_, 50mM NaCl, 1mM MgCl_2_, 0.1 mM CaCl_2_ and 0.01 mM FeCl_3_ supplemented with 0.5% casamino acids and 0.5% glucose](25) in a new microtiter plate, and grown for an additional 24h at 37°C in a humidified chamber. Plates were first measured spectrophotometrically (OD_600_) to determine culture cell density. Supernatants were discarded, and plates were washed 3X with ultrapure water. Plates were dried in a biosafety hood for 2.5h and stained with 200 μL of 0.1% CV for 20 minutes. Plates were then washed 5X to remove excess stain, air-dried for 4h, and 200 μL 30% acetic acid was added to each well. Following a 15 min incubation, CV absorbance was measured spectrophotometrically (OD_560_) and normalized to culture density. Wells exhibiting less than 50% absorbance than the wildtype (MN001) were considered putative hits (134 total) and were subjected to a secondary screen using the same protocol (n=4 for each mutant). Transposon mutants showing a significant reduction in biofilm growth (31 total, as determined by a Mann-Whitney U test) were stored for further characterization.

Biofilm growth of MN001, its Δ*echA* derivative and complemented strain (see below) were also tested using the same microtiter assay. In these experiments, growth medium was supplemented with 2% ethanol to drive expression of the *alcA* promoter in pBMB4 (below). When indicated, *cis*-2-decenoic acid (F13807D, Carbosynth) was also added at a concentration of 310nM (35).

### Arbitrary PCR and mutant sequencing

An arbitrary-PCR-based approach (22) was used to identify sequences flanking transposon insertion sites. The PCR method involved two rounds of reactions, with the first using a primer unique for the mariner transposon and one degenerate primer (pair 1, Table S2)(56). The second round included nested primers (pair 2) unique to the transposon and 5’ end of the arbitrary primer for amplification of PCR products obtained in the first round. PCR products were sequenced at the University of Minnesota Genomics Center (UMGC) and were mapped to the *A. xylosoxidans* MN001 genome (Accession#PRJNA288995).

### Construction and complementation of an *echA* deletion mutant

To generate a deletion vector compatible with *A. xylosoxidans,* pSMV8 (57) was first digested using ApaI. A tetracycline resistance cassette was then amplified from pEX18tc (31) using primer pair 3 (Table S2) and digested with ApaI. Vector and insert (1:3 ratio) were then ligated using T4 ligase, transformed into *E. coli* UQ950 and selected for on LB agar containing tetracycline (15 μg/mL). Colonies were screened using primers M13F and TetR (pair 4) to confirm insertion orientation. One plasmid, pBMB1, was selected for further use.

To generate an in-frame, unmarked deletion of Achromo51_0405 (*echA*), ∼1kb sequences flanking *echA* were PCR amplified using primer pairs 5 and 6 (Table S2). These flanking regions were combined and cloned into pBMB1 digested with SpeI and XhoI using Gibson assembly, resulting in pBMB2 (Figure S1). This plasmid was then chemically transformed into UQ950, and positive ligations were screened by PCR. pBMB2 was then transformed into *E. coli* strain WM3064 and mobilized into *A. xylosoxidans* MN001 by conjugation. Recombinants were selected for on LB-tetracycline agar and double recombinants were selected for on LB agar containing 6% sucrose.

Complementation was achieved via exogenous expression of *echA* (Ax_0405) from pBBR1MCS-5 (57). To do so, an alcohol inducible promoter, *alcA* (32) was first amplified from pGGA008 using primers alcA_F and alcA_R (Table S2) before digestion with restriction enzymes HindIII and BamHI. Ligation into similarly digested pBBR1MCS-5 yielded pBMB3 (pBBR1MCS-5::alcA). *echA* was then amplified from *A. xylosoxidans* MN001 genomic DNA using primers echA_F and echA_R, and digested with BamHI and SacI before ligation into pBMB3 using T4 ligase. The resulting vector, pBMB4 (pBBR1MCS-5::*alcAechA;* Figure S1), was transformed into *E. coli* UQ950. This vector was then introduced into *A. xylosoxidans* MN001 via conjugation with an *E. coli* donor strain WM3064, and transconjugants were selected on LB agar containing 300 μg/mL gentamicin sulfate. All constructs and positive transformants were verified by Sanger sequencing.

### Attachment assay

A modified attachment assay (36) was used to assess early attachment of *A. xylosoxidans* to a polystyrene substratum. Briefly, MN001 and its Δ*echA* derivative were grown for 18h at 37°C in ABT followed by dilution 1:100 into fresh medium. Cells were then grown to mid-log phase (OD_600_ = 0.6) before dilution in ABT to an OD_600_ of 0.1. 200 μl of each culture was added to an 8-chamber coverslip slide (Ibidi, #80824) and incubated at 37°C for 1h. Following incubation, slides were rinsed twice with 200 μl of PBS to remove unattached biomass, and attached cells were stained using SYTO 9 (Invitrogen) in PBS according the manufacturer’s protocol. Substrata were imaged using an Olympus IX83 inverted fluorescence microscope with a transmitted Koehler illuminator and a 40X objective lens (Olympus). Four images per strain per biological replicate (n=4) were captured on a Hamamatsu ORCA camera, and post-acquisition analysis was performed using FIJI software (58) by calculating the integrated density of SYTO 9.

### Colony biofilm assay

MN001 and Δ*echA* were grown overnight in LB medium, diluted 1:1000, and 10μL was spotted on nutrient agar containing 1% tryptone, 1% agar, 20μg/mL Coomassie Brilliant Blue, and 40 μg/mL Congo red (36). Plates were incubated at 37°C for 6 days and monitored daily for colony morphology.

### Scanning electron microscopy

Overnight cultures were diluted 1:10 in fresh LB medium and were added to 48-well microtiter plates containing autoclaved Aclar fluoropolymer film (Electron Microscopy Sciences, Hatfield, PA). Biofilms were grown for 48 h at 37°C, shaking at 50 rpm, and prepared for SEM using cationic dye stabilization methods (59,60). Briefly, Aclar membranes containing biofilm growth were washed three times in 0.2M sodium cacodylate buffer, and submerged in primary fixative (0.15M sodium cacodylate buffer, pH 7.4, 2% paraformaldehyde, 2% glutaraldehyde, 4% sucrose, 0.15% alcian blue 8 GX) for 22h. Samples were washed three more times prior to a 90 minute treatment with secondary fixative (1% osmium tetroxide, 1.5% potassium ferrocyanide, 0.135M sodium cacodylate, pH 7.4). After three final washes, biofilms were chemically dehydrated in a graded ethanol series (25, 50, 70, 85, 95 [2x] and 100% [2x]) before CO_2_-based critical point drying. Aclar membranes were attached to SEM specimen mounts using carbon conductive adhesive tape and sputter coated with ∼5nm iridium using the Leica ACE 600 magnetron-based system. Biofilms were imaged using a Hitachi S-4700 field emission SEM with an operating voltage of 2kV.

### Antibiotic challenge

Biofilm antimicrobial susceptibility testing was performed using a chamber slide assay (61). Briefly, MN001 and Δ*echA* were grown in LB overnight, diluted 1:1000 and grown to an OD_600_ of 0.6 before dilution to an OD_600_ of 0.5. 200μL of each culture was then added to each well of an Ibidi 8-chamber coverslip slide and incubated at 37°C in a humidified chamber. After 24h, medium was replaced with fresh LB and incubated for an additional 24h. Media was gently aspirated from each well, replaced with LB containing either tobramycin or levofloxacin (0, 10, 100, and 1000 μg/mL), and incubated at 37°C for 6h. Cells were washed in sterile PBS to remove unattached biomass, stained for 15 minutes using the BacLight Live/Dead viability assay (Life Technologies, #L7012) and visualized by fluorescent imaging as described above. Integrated density for SYTO 9 and propidium iodide (PI) for each image was determined using FIJI (58), and percentage of dead biomass was determined by (average integrative density of PI)/(average integrated density of PI + integrated density of SYTO 9)(61). Data were generated using four biological replicates (n=4).

## ACKNOWLEDGEMENTS

pGGA008 (Addgene plasmid #48817) was a gift from Dr. Jan Lohmann (Heidelberg). We thank Clayton Summitt for assistance with data analysis and other members of the Hunter lab for critical review of the manuscript. Parts of this work were carried out in the Characterization Facility, University of Minnesota, which receives partial support from NSF through the Materials Research Science and Engineering Centers (MRSEC) program. This study was supported by the University of Minnesota, a NHLBI Pathway to Independence award to RCH, American Society for Microbiology Undergraduate Research Fellowships to AKL and LAK.

## AUTHOR CONTRIBUTIONS

L.C. performed experiments, created figures, and helped write and edit the manuscript. B.B. performed experiments, created figures, and helped edit the manuscript. C.P. and A.L. performed experiments. L.K. performed SEM experiments and created figures. R.H. contributed to the conception, experimental design, data interpretation, statistical analyses, and manuscript preparation.

## DATA AVAILABILITY

All data generated during and/or analyzed during the current study are included in this article, its supplementary information files, or are available from the corresponding author on reasonable request.

