## Supplemental Information for "A putative enoyl-CoA hydratase contributes to biofilm formation and the antibiotic tolerance of *Achromobacter xylosoxidans*"

**Cameron et al.**

### SUPPLEMENTARY INFORMATION

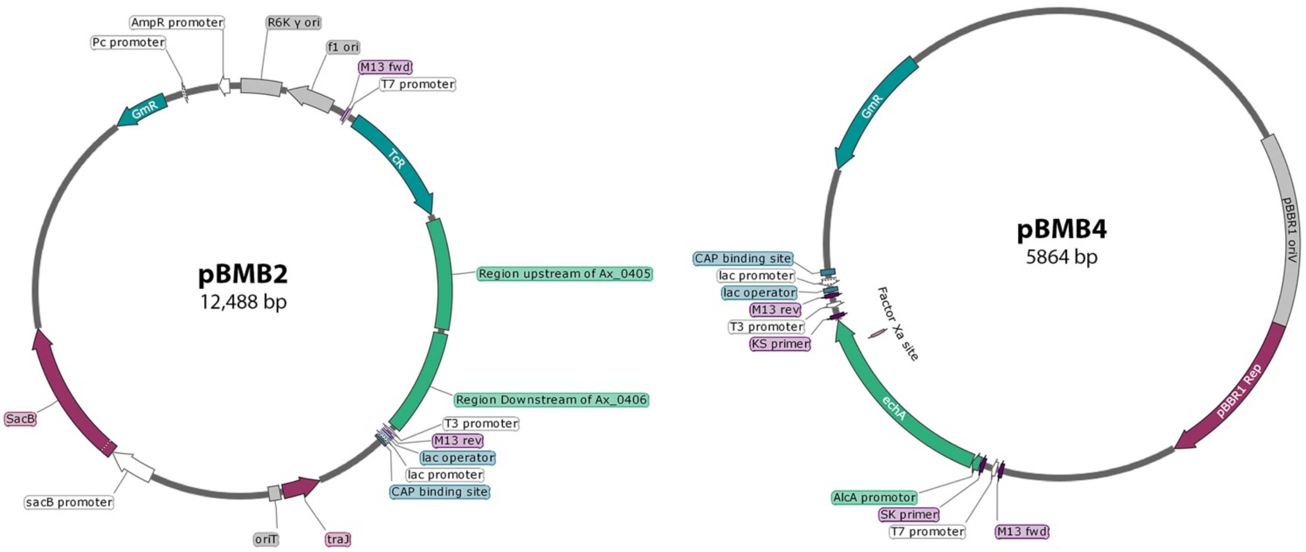

**Figure S1.** Deletion and complementation constructs.

**Table S1.** Strains and plasmids used in this study

| **Strain or Plasmid** | **Characteristic** | **Source** |
|---|---|---|
| *A. xylosoxidans* | | |
| MN001 | Cystic fibrosis clinical isolate | (1) |
| Δ*echA* | MN001 with a deletion of Axylo_0405 (*echA*) | This study |
| *E. coli* | | |
| β2155 | Donor strain for conjugations | (2) |
| WM3064 | Donor strain for conjugations | (3) |
| UQ950 | DH5α λpir | (3) |
| Plasmids | | |
| pTnTet | pSC123 with $Tet^R$ cassette from pBSL199 | (4) |
| pSMV8 | Mobilizable suicide vector, $Gm^R$ | (3) |
| pBMB1 | pSMV8 with $Tet^R$ cassette from pEX18tc | This study |
| pEX18tc | Gene replacement vector with MCS from pUC18 | (5) |
| pBMB2 | Mobilizable suicide vector for *echA* deletion, $Tet^R$ | This study |
| pBBR1MCS-5 | Complementation vector | (6) |
| pGGA008 | Cloning vector containing *alcA* promoter, $Amp^R$ | (7) |
| pBMB3 | Complementation vector pBBR1MCS-5 with an alcohol inducible (alcA) promoter | This study |
| pBMB4 | Complementation vector pBBR1MCS-5::*alcAechA* | This study |

**Table S2.** Primers used in this study.

| Primer Pair | Name | Sequence (5'-3') |
|---|---|---|
| 1 | ARB1 | GGCCACGCGTCGACTAGTACNNNNNNNNNNGATAT |
| | TnTet1 | AACAAGCCAGGGATGTAACG |
| 2 | ARB2 | GGCCACGCGTCGACTAGTAC |
| | TnTet2 | TGTCAGACCGGGGACTTATC |
| 3 | TetF | NNNGGGCCC*GCTAGCTTTAATGCGGTAGT* |
| | TetR | NNNNGGGCCCTGGAGTGGTGAATCCGTTAG |
| 4 | M13F | TGTAAAACGACGGCCAGT |
| | TetR | NNNNGGGCCCTGGAGTGGTGAATCCGTTAG |
| 5 | *echA* upF | ccctcgaggtcgacggtatcgataTTCAGGGTCAGTTCGCTCAT |
| | *echA* upR | TGTTCCAGCGTGATATCGGT |
| 6 | *echA* downF | ACCGATATCACGCTGGAACAGCTCATGCGAAGGTCCTGG |
| | *echA* downR | cggtggcggccgctctagaactagtACACGTGACGCCGTTATGC |
| 7 | *alcA*_F | cgatAAGCTTcgggatagttccgacctaggatt |
| | *alcA*_R | cgatGGATCCttatatagatgttcagctatgcg |
| 8 | *echA*_F | atcgGGATCCatgaccgatatcacgctgg |
| | *echA*_R | cgatGAGCTCtcagaagggcgagtcgg |
